# Protein-carbohydrate ingestion alters Vps34 cellular localization independent of changes in kinase activity in human skeletal muscle

**DOI:** 10.1101/2020.02.10.941054

**Authors:** Nathan Hodson, Jessica R. Dent, Zhe Song, Mary F. O’Leary, Thomas Nicholson, Simon W. Jones, James T. Murray, Stewart Jeromson, D. Lee Hamilton, Leigh Breen, Andrew Philp

## Abstract

The mechanistic target of rapamycin (mTOR) complex 1 (mTORC1) regulates cell size and growth in response to nutrients, however, the mechanisms by which nutrient levels are sensed by mTORC1 in human skeletal muscle are yet to be fully elucidated. The Class III PI3Kinase Vps34 has recently been proposed as a sensor essential for mTORC1 activation following nutrient stimulation. We therefore investigated the effects of increasing nutrient availability through protein-carbohydrate (PRO-CHO) feeding on Vps34 kinase activity and cellular localization in human skeletal muscle. Eight young, healthy males (age – 21 ± 0.5yrs, mean ± SEM) ingested a PRO-CHO beverage containing 20/44/1g PRO/CHO/FAT respectively, with skeletal muscle biopsies obtained at baseline and 1h and 3h post-feeding. PRO-CHO feeding did not alter Vps34 kinase activity, but did stimulate Vps34 translocation toward the cell periphery (PRE (mean±SEM) - 0.273±0.021, 1h - 0.347±0.022, Pearson’s Coefficient (r)) where it co-localized with mTOR (PRE – 0.312±0.018, 1h – 0.348±0.024, Pearson’s Coefficient (r))). These alterations occurred in parallel to an increase in S6K1 kinase activity – 941±164% of PRE at 1h post-feeding). Subsequent *in vitro* experiments in C2C12 and human primary myotubes displayed no effect of the Vps34-specific inhibitor SAR405 on mTORC1 signalling responses to elevated nutrient availability. Therefore, in summary, PRO-CHO ingestion does not increase Vps34 activity in human skeletal muscle, whilst pharmacological inhibition of Vps34 does not prevent nutrient stimulation of mTORC1 *in vitro*. However, PRO-CHO ingestion promotes Vps34 translocation to the cell periphery, enabling Vps34 to associate with mTOR. Therefore, our data suggests that interaction between Vps34 and mTOR, rather than changes in Vps34 activity *per se* may be involved in PRO-CHO activation of mTORC1 in human skeletal muscle.

## Introduction

Amino acids (AAs) are critical to skeletal muscle plasticity, acting as both substrates in the process of muscle protein synthesis (MPS) as well as initiating the signaling pathways which activate this cellular process (32, 33). Carbohydrate (CHO) ingestion can also elevate MPS via insulin action (3), and a combination of these nutrients is believed to act synergistically on MPS following exercise (20). In skeletal muscle, it is believed that increases in MPS are governed primarily by the activation of the mechanistic target of rapamycin complex 1 (mTORC1) (5, 6), an evolutionarily conserved serine/threonine kinase complex which stimulates translation initiation and elongation (14, 16, 31) in response to increased nutrient provision.

The canonical mechanism by which AAs stimulate mTORC1 activity is thought to be through the elevation of mTORC1 complex co-localization with the lysosome (27) *in vitro*, or through mTORC1/lysosomal trafficking *in vivo/vitro* (11, 15, 28). However, how nutrients stimulate mTORC1 activity in human skeletal muscle is still poorly understood. A potential nutrient-sensitive activator of mTORC1 is the vacuolar protein sorting 34 (Vps34), a class III PI3Kinase. The primary function of Vps34 is the production of phosphatidylinositol 3-phosphate (PI(3)P) through the phosphorylation of phosphatidylinositol (2), a product responsible for the recruitment of various proteins to phospholipid bilayers (i.e. plasma and lysosomal membranes) (8). A role for Vps34 in nutrient sensing was first proposed by Byfield et al. (4), who reported that overexpression of Vps34 in HEK293 cells elicited a 2-fold increase in S6K1 activity, a common readout of mTORC1 activation. Conversely, siRNA targeting Vps34 abolished insulin-stimulated S6K1^Thr389^ phosphorylation (4). Nobukuni et al. (22) reiterated these findings, displaying that siRNA-mediated reductions in Vps34 expression, in HEK293 cells, dramatically attenuated mTORC1 activation in response to both AA and insulin stimulation. In addition, recent *in vitro* evidence suggests that Vps34 colocalises with mTOR, close to cellular membranes, following insulin stimulation (10), and is required for nutrient-stimulated translocation and activation of mTORC1 (10). As such, Vps34 represents a novel candidate as a nutrient-sensitive activator of mTORC1.

With regard to skeletal muscle, 3h and 24h exposure to leucine (5mM) and insulin (100nM) elevated Vps34 protein content and mTOR^Ser2448^ and S6K1^Thr389^ phosphorylation in human primary myotubes (9), whilst supra-physiological levels of AA’s increases Vps34 activity in C2C12 myotubes (17). In addition, high frequency electrical contraction has been reported to increase Vps34 activity in rodent *Tibialis Anterior* muscle (17), whereas sprint exercise and protein ingestion failed to activate Vps34 in human skeletal muscle (26). Overall, such data implicates a possible role for Vps34 in nutrient/contraction sensing within skeletal muscle. However, a more detailed investigation in human skeletal muscle is required.

Therefore, our primary aim was to investigate if AA/CHO feeding could affect Vps34 activity and cellular localisation in human skeletal muscle. We hypothesised that Vps34 activity would increase in response to AA/CHO feeding in parallel to increases in mTORC1 signaling. Our secondary aim was to examine whether inhibition of Vps34 kinase activity *in vitro* with the specific inhibitor SAR405 (24, 25) would attenuate nutrient-activation of mTORC1.

## Methods

### Participants

Eight young, healthy males (age – 21 ± 0.5yrs, mean ± SEM) volunteered to partake in the current study. All participants were considered healthy (as assessed by a general health questionnaire) and recreationally active (∼3 exercise sessions per week) but not involved in a structured exercise training program. Exclusion criteria encompassed current cigarette smokers, recreational drug users (including anabolic steroids), the presence of neuromuscular disease and any medication/condition that may affect nutrient digestion/absorption i.e. inflammatory bowel disease. Participants provided written informed consent prior to participation and all procedures were approved by the NHS West Midlands Black Country Research Ethics Committee (15/WM/0003) and conformed to the standards set out in the Declaration of Helsinki (7^th^ version).

### Study Design

On the day of the experimental trial, participants reported to the laboratory following an overnight fast (∼10h) and having refrained from strenuous exercise and alcohol consumption in the prior 48h. Upon arrival, participants were placed in a supine position and a 21G cannula was inserted into the antecubital vein of one arm to allow for repeated blood sampling. At this point an initial baseline blood sample was obtained from all participants. A skeletal muscle biopsy sample was then taken from the *vastus lateralis* of a randomised leg using the Bergstrom percutaneous needle technique, modified for suction (30). Participants then consumed a commercially available protein-carbohydrate beverage (Gatorade Recover®, Gatorade, Chicago, IL, USA.) providing 20/44/1g of protein, carbohydrate and fat respectively. Further venous blood samples were taken every 20 minutes for a 3h post-prandial period and subsequent skeletal muscle biopsy samples were obtained at 1h and 3h following beverage ingestion. Muscle samples were blotted free of excess blood and dissected free of any excess adipose and connective tissue, then immediately frozen in liquid nitrogen and stored at -80°C until analysis. A separate piece of muscle tissue was placed in optimal cutting temperature (OCT) compound (VWR, Lutterworth, UK.) and frozen in liquid nitrogen-cooled isopentane before storage at - 80°C. Blood samples were collected into EDTA-coated vacutainers (BD, Franklin Lakes, NJ, USA.) and then centrifuged at 1000g for 15min to separate plasma. Plasma was then aliquotted into micro-centrifuge tubes and stored at -80°C until analysis. The experimental design is depicted in Figure 1.

**Figure 1.**
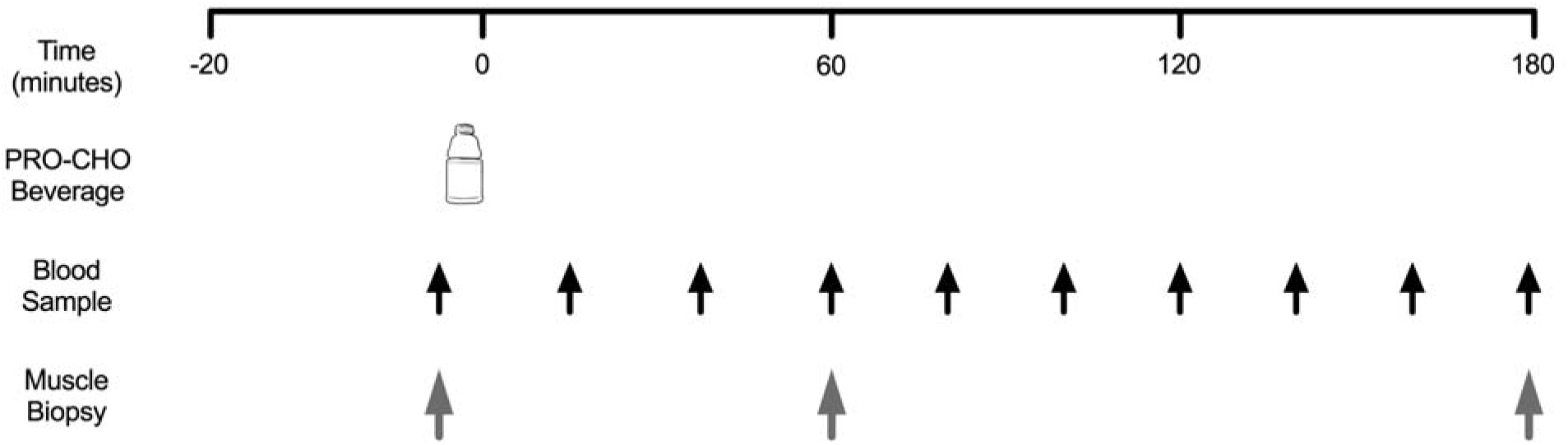
Schematic of Experimental Protocol for Human Trial.

### Blood analyses

Plasma insulin concentrations were quantified using a commercially-available ELISA kit (IBL International, Hamburg, Germany.) as per the manufacturer’s instructions. Plasma leucine concentrations were determined via gas chromatography-mass spectrometry (GC-MS) using an internal standard method, as previously described (19), following the conversion of plasma free amino acids to their N-tert-butyldimethyl-silyl-N-methyltrifluoracetamide (MTBSTFA) derivative.

### S6K1 and AKT Kinase Activity Assays

S6K1 and AKT kinase activity assays were conducted as described previously (18) with the following antibodies; S6K1 – SCBT no.2708 (Santa Cruz Biotechnologies, Dallas, TX, USA.) & AKT (DSTT, Dundee, UK). Briefly, a ∼30mg piece of muscle tissue was homogenized on ice in RIPA buffer (50 mmol/l Tris·HCl pH 7.5, 50 mmol/l NaF, 500 mmol/l NaCl, 1 mmol/l sodium vanadate, 1 mmol/l EDTA, 1% (vol/vol) Triton X-100, 5 mmol/l sodium pyrophosphate, 0.27 mmol/l sucrose, and 0.1% (vol/vol) 2-mercaptoethanol and Complete protease inhibitor cocktail (Roche)). Cellular debris was then removed via centrifugation at 13000g for 15min (4°C). Protein concentrations of samples was then determined via bicinchoninic acid (BCA) protein assay. Immunoprecipitation of the target protein was then conducted on 200µg protein for 2h at 4°C in homogenization buffer (50 mM Tris·HCl pH 7.5, 0.1 mM EGTA, 1 mM EDTA, 1% (vol/vol) Triton X-100, 50 mM NaF, 5 mM NaPPi, 0.27 M sucrose, 0.1% -mercaptoethanol, 1 mM Na3(OV)4, and 1 Complete (Roche) protease inhibitor tablet per 10 mL) combined with 2.5µL Protein G Sepharose beads and appropriate antibody. Immunoprecipitates were subsequently washed twice in high-salt buffer (homogenization buffer with 0.5M NaCl added) and once in assay buffer (50 mM Tris·HCl pH 7.4, 0.03% Brij35, and 0.1% -mercaptoethanol). Immunoprecipitates were then resuspended in 10µL assay buffer and activity assay commenced every 20 seconds through the addition of a hot assay mix (assay buffer + 100µM ATP + 10mM MgCl_2_ + 32γATP + synthetic substrate (S6tide - KRRRLASLR at 30 µM & Crosstide - GRPRTSSFAEG at 30µM for S6K1 and AKT assays respectively). Every 20s reactions were stopped through spotting on to chromatography paper, immersion in 75mM phosphoric acid and drying. Chromatography paper was immersed in GoldStar LT Quinta Scintillation fluid (Meridian Biotechnologies, Chestefield, UK) and spots were counted in a Packard 2200CA TriCarb Scintillation Counter (United Technologies) as fmol·min^-1^·mg^-1^.

### Vps34 Kinase Activity Assay

Vps34 kinase activity assays were conducted as previously described (17) from 25mg muscle homogenized in Cantley lysis buffer. Vps34 was immunoprecipitated overnight at 4°C from tissue lysates containing ∼1mg total protein using 2µg anti-Vps34 antibody (sheep antibody produced by Dr. James T. Murray, Trinity College Dublin) before immobilisation on Protein G Sepharose beads for 1h. Immunoprecipitates were then washed 3 times in Cantley lysis buffer, once in Tris-LiCl (10 mM Tris, pH 7.5, 5 mM LiCl, 0.1 mM Na2VO4) and twice in TNE (10 mM Tris, pH 7.5, 150 mM NaCl, 1 mM EDTA, 0.1 mM Na2VO4) and then resuspended in 60 μl TNE+ (TNE, 0.5 mM EGTA, pH 8.0, 1 : 1000 2-mercaptoethanol). Samples were then incubated with 20µg Vps34 antigen peptide for 10min before 10μl of 30 mM MnCl2 and 10μl of 2 mg ml−1 phosphoinositol were added to provide substrate for the reaction. Reactions then commenced through the addition of 5µL assay buffer (400 μM unlabelled ATP, 12.5 μCi of 32γATP, 4.3 μl water) for 10 minutes at 30°C. Reactions were terminated by the addition of 20µL 8M HCl, phase separated using 1:1 chloroform and methanol and the lower phase spotted onto an aluminium-backed 60 A silica ° TLC plate (Merck, Damstadt, Germany). This was then run in a TLC chamber solvent system to determine 32γP transfer to substrate.

### Immunohistochemistry

Immunohistochemical analysis was conducted as described previously (28). In short, 5µm sections of muscle tissue were sectioned at -25°C using a Bright 5040 Cryostat (Bright Instrument Company Ltd., Huntingdon, UK) and transferred to room temperature (RT) glass slides (VWR international, UK) and allowed to airdry for ∼1h. Sections from each time point for each participant were sectioned onto the same slide in duplicate to remove slide-to-slide variation during analysis. Muscle sections were subsequently fixed in a 3:1 solution of acetone and ethanol, washed 3 times in Phosphate Buffered Saline (PBS) before incubation in relevant primary antibodies (antibodies and dilutions in Table 1) diluted in 5%NGS to prevent non-specific secondary binding for 2h at RT. Subsequently, sections were again washed in PBS and then incubated in corresponding secondary antibodies (details in Table 1) for 1h at RT. Following further washes, slides were then incubated in Wheat Germ Agglutinin (WGA – conjugated to 350nm fluorophore) for 30min at RT in order to mark the sarcolemmal membrane. After a final wash in PBS, slides were then mounted in Mowiol® 4–88 (Sigma-Aldrich, Poole, UK) to protect fluorophores and a glass coverslip was applied. Slides were then left to dry overnight in a dark cabinet prior to image capture. Pilot stains were also conducted with and without the presence of the Vps34 primary antibody to ensure no non-specific binding of the secondary antibody.

**Table 1.** Summary of Antibodies Used.

| Primary antibody | Source | Dilution | Secondary antibody | Dilution |
| --- | --- | --- | --- | --- |
| Monoclonal anti-mTOR antibody with mouse antigen, isotype IgG γ1 kappa | Millipore, 05-1592 | 1:200 | Goat anti-mouse IgG γ1 kappa Alexa@594 | 1:200 |
| Polyclonal anti-Lamp2 antibody with rabbit antigen, isotype IgG | Abgent, AP1824d | 1:100 | Goat anti-rabbit IgG(H+L) kappa Alexa@488 | 1:200 |
| Monoclonal anti-Vps34/PIK3C3 antibody with rabbit antigen, isotype IgG | Cell Signaling Technology #3358 | 1:20 | Goat anti-mouse IgM Alexa@594 | 1:200 |
| Wheat Germ Agglutinin-350 | W11263, Invitrogen | 1:20 | Alexa Fluor® 350 Conjugated | N/A |

### Image Capture and Analysis

Stained muscle sections were observed under a Nikon E600 widefield microscope using a 40×0.75NA objective under three colour filters achieved by a SPOT RT KE colour three shot CCD camera (Diagnostic Instruments Inc., MI, USA) illuminated by a 170W Xenon light source. In the current study, DAPI UV (340-380nm) excitation filter was utilized to visualize WGA, TxRed (540-580nm) for mTOR visualization and FITC (465-495nm) for LAMP2.Vps34 depending on the stain conducted. For each time point, approximately 8 images were taken per section, each consisting of ∼8 muscle fibers. As sections were analysed in duplicate, approximately 120 muscle fibres per time point per participant were included in analysis. Image processing and quantification was completed on ImageProPlus 5.1 software (Media Cybernetics, MD, USA.) with all variables kept consistent for all sections on a given slide. Prior to co-localisation analysis, all images underwent a no neighbour deconvolution algorithm as a filter. Pearson’s correlation coefficient (Image-Pro software) was used to quantify co-localization of proteins stained in different channels. This method of assessing co-localization was utilized as it measures co-localization on a pixel-by-pixel basis and is relatively free of user bias (7).

### In vitro experiments

C2C12 myoblasts were purchased from American Type Culture Collection (ATCC, Manassas, VA, USA.) and cultured on 150mm culture plates in high glucose Dulbecco’s minimum essential medium (DMEM, ThermoFisher Scientific, Waltham, MA, USA.) supplemented with 10% foetal bovine serum (FBS, Hyclone, VWR, Lutterworth, UK.) and 1% penicillin-streptomycin (PS, ThermoFisher Scientific). When 80% confluent, cells were trypsinized (0.05% Trypsin-EDTA, ThermoFisher Scientific) and seeded onto 6-well plates at a density of 2×10^5^cells/well. Myoblasts were then cultured until ∼95% confluency (∼36h) at which time media was changed to elicit differentiation of myoblasts to myotubes (DMEM supplemented with 2% horse serum (HS, Hyclone, VWR) and 1% PS). Differentiation was allowed to occur for 5 days, with media replaced every other day, until myotubes were fully formed.

At this point, myotubes were nutrient deprived in Earl’s Balanced Salt Solution (EBSS, ThermoFisher Scientific) for ∼14h, with a subset of myotubes maintained in DMEM (2%HS, 1%PS) to serve as a ‘baseline’ condition. Following nutrient deprivation, a subset of myotubes were collected and the remaining myotubes were split into 2 conditions, serum recovery and serum recovery + Vps34 inhibition. Vps34 inhibition was achieved via the addition of the specific Vps34 inhibitor SAR405 (10µM) for 1h prior to serum recovery, a concentration and incubation time previously shown to fully inhibit Vps34 kinase activity *in vitro* (25). Serum recovery occurred through the removal of EBSS and addition of DMEM (2%HS) for 30min prior to collection. Before collection, myotubes were washed twice in ice-cold PBS, before being scraped into 150 µL ice-cold sucrose lysis buffer (50 mM Tris, 1 mM EDTA, 1 mM EGTA, 50 mM NaF, 5 mM Na_4_P_2_O_7_-10H_2_O, 270 mM sucrose, 1 M Triton-X, 25 mM β-glycerophosphate, 1µM Trichostatin A, 10 mM Nicatinamide, 1 mM 1,4-Dithiothreitol, 1% Phosphatase Inhibitor Cocktail 2 (Sigma), 1% Phosphatase Inhibitor Cocktail 2 (Sigma), 4.8% cOmplete Mini Protease Inhibitor Cocktail; (Roche). Lysates were then immediately frozen in liquid nitrogen and stored at -80°C until analysis. Experiments were conducted in triplicate at 3 separate passage numbers equalling n=9 for statistical analysis.

Human primary myoblasts were isolated from 4 patients (age 61±6yrs, BMI 28.7±0.65kg/m^2^, mean±SEM) as previously described (23). Cells were passaged at 60% confluency on 0.2% gelatin-coated 100mm culture plates in Hams F10 media (ThermoFisher Scientific, supplemented with 20% FBS and 1% PS) to prevent spontaneous fusion of myoblasts to myotubes, and at passage 3 were seeded onto 6-well plates at a density of 5×10^4^ cells/well. Myoblasts were then cultured to 80-90% confluency, at which time media was changed to induce differentiation to myotubes (F10 supplemented with 6% HS and 1% PS). Once myotubes were fully formed (6-10days), experiments were conducted in a similar fashion to those described above for C2C12 myotubes with certain alterations. Baseline conditions for human primary myotubes were ∼14h incubation in Hams F10 media (20%FBS, 1%PS) and serum recovery experiments were conducted for 30min in Hams F10 (20%FBS, 1%PS) following ∼14h EBSS incubation. All other experimental variables were consistent between C2C12 and human primary experiments and cells were collected in an identical fashion. Experiments were run in triplicate for myotubes isolated from each patient and the mean of these results utilized for statistical analysis.

Cell lysates were subsequently homogenised by sonication (3×15s at 50% maximal wattage) and centrifuged at 8000g for 10mins at 40°C to remove insoluble material. Protein content of these lysates was then determined by DC protein assay (BioRad, Hercules, CA, USA.) and samples were diluted to a desired protein concentration in 1x Laemmli sample buffer and boiled at 95°C for 5 minutes to denature proteins.

### Immunoblotting

Immunoblotting analysis was conducted as described previously (29). Briefly, equal amounts of protein were loaded into 8-15% polyacrylamide gels and separated by SDS-PAGE. Proteins were then transferred to BioTrace NT nitrocellulose membranes (Pall Life Sciences, Pensacola, FL, USA.) and stained with Ponceau S as a loading control. Membranes were then blocked in 3% skimmed-milk diluted in Tris-buffered Saline with tween (TBST) for 1h at RT. Following washing in TBST, membranes were then incubated overnight in relevant primary antibodies, subsequently washed again and incubated in corresponding HRP-conjugated secondary antibodies (anti-rabbit IgG #7074, Cell Signaling Technologies (CST), Danvers, MA, USA. 1:10000). Enhanced chemiluminescence HRP detection kit (Merck-Millipore, Watford, UK.) was used to quantify antibody binding. Each phosphorylated protein visualized was expressed in relation to its total protein content, after each target had been normalized to a loaded control (Ponceau). All primary antibodies utilized for immunoblotting were purchased from CST and diluted at 1:1000 in TBST unless stated otherwise: p70 ribosomal S6 kinase 1 (S6K1, #2708), p-S6K1^Thr389^ (#9205), ribosomal protein S6 (S6, #2217), p-S6^Ser235/236^ (#4858), p-S6^Ser240/244^ (#5364), eukaryotic translation initiation factor 4E-binding protein 1 (4EBP1, #9452, 1:500) and p-4EBP1^Thr37/46^ (#9459).

### Statistical analysis

Alterations in enzyme kinase activity, protein-protein colocalization, plasma insulin and plasma leucine concentrations were analysed utilizing a repeated measures analysis of variance (ANOVA) with one within-subject factor (time). Changes in phosphorylation status of proteins in human primary myotubes was also analysed with a repeated measures ANOVA with one within-subject factor (condition). A one-way ANOVA with one between-subject factor (condition) was used to analyse changes in phosphorylation status of proteins in C2C12 myotubes. Greenhouse-Geisser corrections were applied to F values if data did not pass Mauchly’s test of sphericity. If a significant main effect was found, *post-hoc* analysis was conducted on comparisons determined *a priori* with the Holm-Bonferroni correction for multiple comparisons. Significance for all variables was set at p<0.05 and data are presented as mean±SEM unless otherwise stated.

## Results

### Blood Analyses

For plasma insulin analysis, physiological results from 7 out of 8 participants were obtained and, as such, statistical analysis was completed on n=7. A significant time effect was observed for changes in plasma insulin concentrations (p<0.001), however, following *post hoc* analysis no differences between individual time points were noted (p>0.05, Figure 2A). A significant time effect was also observed for plasma leucine concentrations (p<0.001). Plasma leucine concentrations were elevated above basal levels at 20min post-feeding (0.113±0.006 vs. 0.187±0.016mmol/L, p=0.02, Figure 2B) and remained above baseline (all p<0.012) until 160min post-feeding (0.113±0.006 vs. 0.118±0.006, p=0.51).

**Figure 2.**
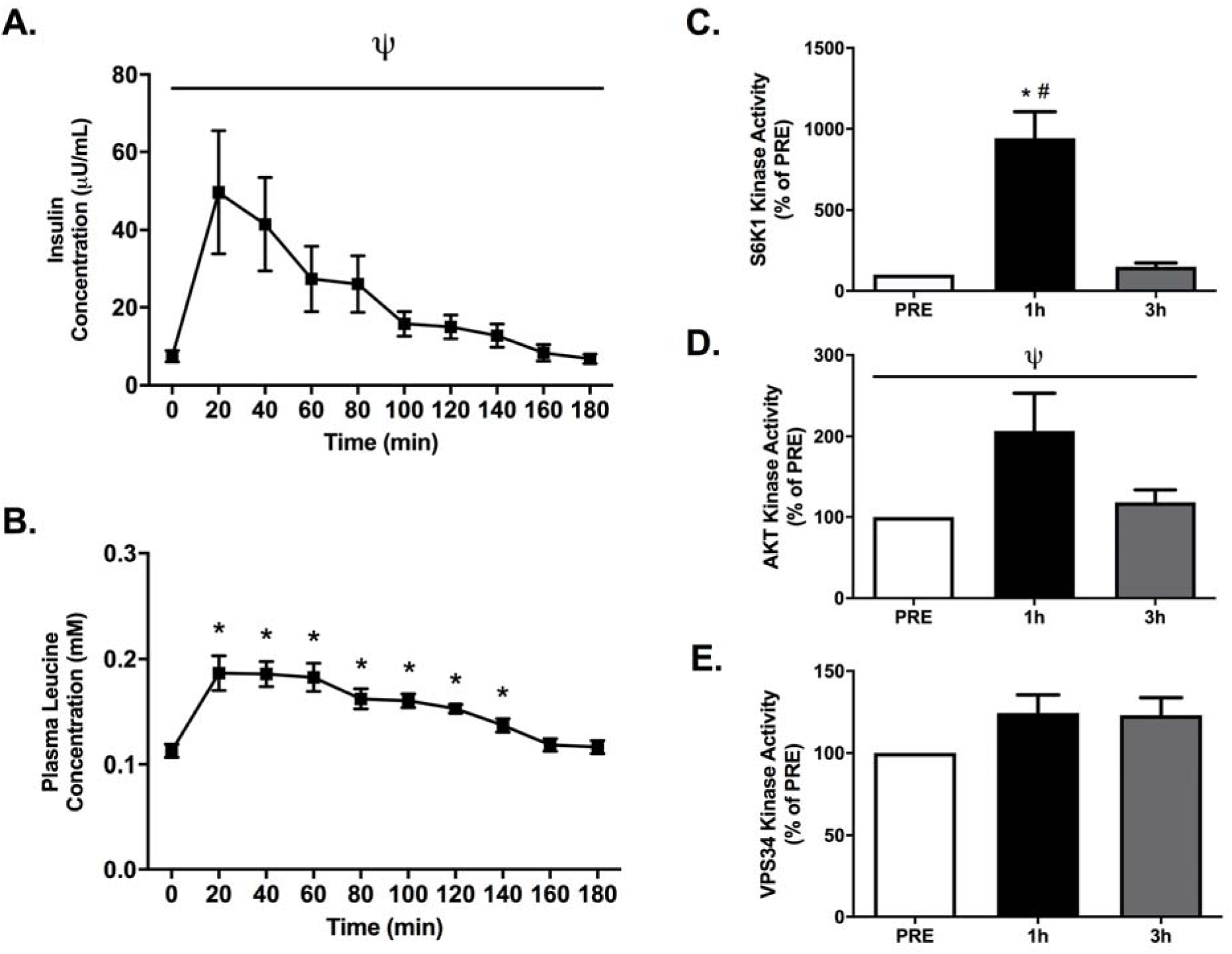
The effect of protein-carbohydrate feeding on plasma insulin and leucine concentrations and enzyme kinase activities. Insulin concentrations (A) are presented as µU/ml and Leucine concentrations (B) are presented as mM. Kinase activity of S6K1 (C), AKT (D) and Vps34 (E) are presented as % of PRE. For A & B, Ψ denotes a significant effect of time (p<0.05) and *denotes a significant difference at this time point compared to 0 (p<0.05). For C, D & E *denotes a significant difference at this time point compared to PRE (p<0.05), ^#^denotes a significant difference at this time point compared to 3h (p<0.05) and Ψ denotes a significant effect of time (p<0.05). All values are presented as mean±SEM. Data analyzed on SPSS using Repeated Measures ANOVA with Holm-Bonferroni *post hoc* comparisons conducted on Microsoft Excel. Insulin – n=7, Leucine – n=8. Kinase acitivies – n=8

### Kinase Activity Assays

A significant time effect was observed for S6K1 activity (p=0.001), with S6K1 activity significantly higher 1h post-feeding compared to PRE and 3h post-feeding (1h – 941.3±164.6% of PRE, 3h – 149.6±23.2% of PRE, p=0.003 & p=0.004 respectively, Figure 2C). Activity of S6K1 also trended toward being greater at 3h post-feeding compared to PRE (p=0.07, Figure 2C). A significant time effect was also apparent for AKT activity (p=0.05, Figure 2D). Following *post hoc* analysis, however, no differences in AKT activity between individual time points was apparent (1h – 206.9±46.6% of PRE, 3h – 118.1±15.5% of PRE, p>0.05, Figure 2D). Finally, no differences in Vps34 activity were noted at any time point (p>0.05, Figure 2E).

### Co-localization

No time effect for mTOR co-localization with LAMP2 (lysosomal marker) was found (p=0.347, Figure 3B) suggesting these proteins are co-localized independently of a nutritional stimulus. A significant time effect was observed for mTOR co-localization with WGA (membrane marker, p=0.026). Following feeding, mTOR-WGA co-localization increased by 17% at 1h before returning to basal values by 3h, however, following *post hoc* analysis, no alterations were significant (PRE – 0.186±0.009, 1h – 0.212±0.014, 3h – 0.184±0.009, PRE vs. 1h p=0.090, 1h vs. 3h p=0.067, Figure 3C). Vps34 co-localization with WGA exhibited a trend toward a time effect (p=0.053) and subsequent *post hoc* analysis revealed that co-localization was greater 1h post-feeding compared to PRE feeding levels (0.347±0.022 vs. 0.273±0.021, p=0.043, Figure 4B), however no other differences were apparent (p>0.05). Finally, there was a significant effect of time observed for mTOR co-localization with Vps34 (p=0.045). Here, following *post hoc* analysis no differences between individual time points were apparent (p>0.05), although a trend toward a greater mTOR-Vps34 co-localization 1h post-feeding compared to 3h was noted (0.347±0.024 vs. 0.315±0.016, p=0.067, Figure 4C).

**Figure 3.**
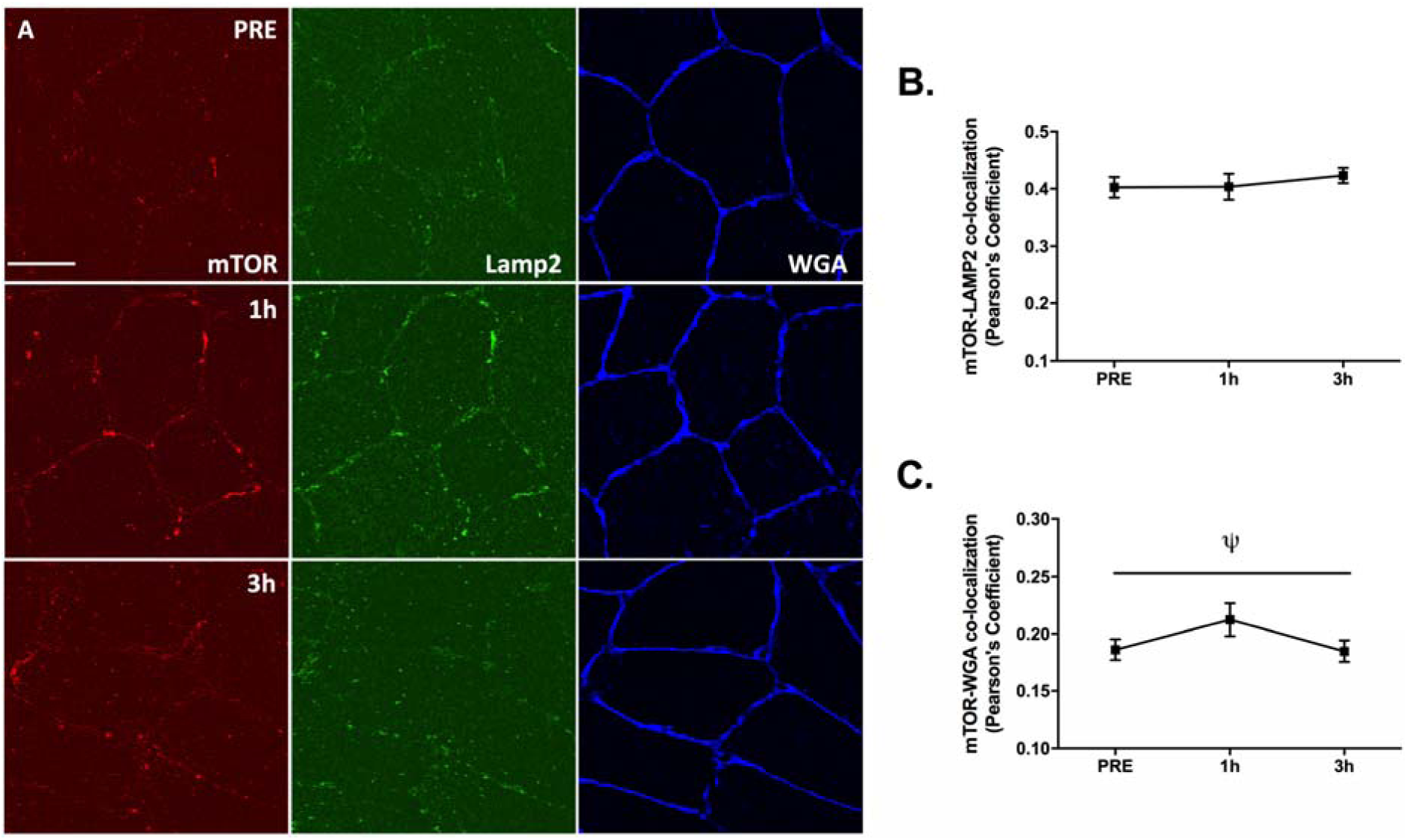
The effect of protein-carbohydrate ingestion on mTOR-LAMP2 and mTOR-WGA co-localization. Representative images of mTOR (red), LAMP2 (green) and WGA (blue) stains at each time point are provided (A). Quantification of mTOR-LAMP2 (B) and mTOR-WGA (C) co-localization is presented as Pearson’s correlation coefficient. Data in B and C are presented as mean±SEM. Ψ denotes a significant effect of time (p<0.05). Data analyzed on SPSS using Repeated Measures ANOVA with Holm-Bonferroni *post hoc* comparisons conducted on Microsoft Excel. All analyses – n=8.

**Figure 4.**
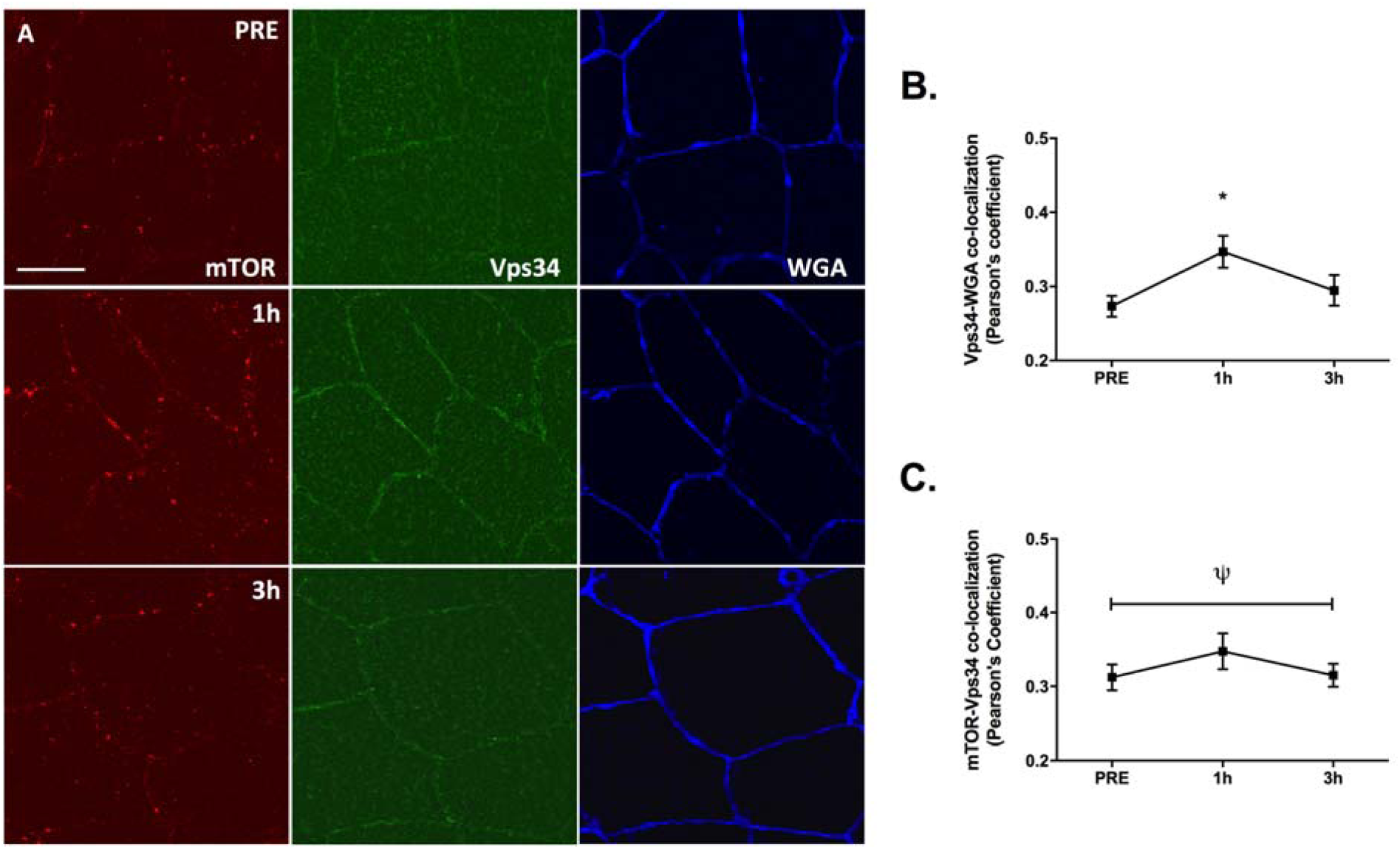
The effect of protein-carbohydrate ingestion on mTOR-VPS34 and mTOR-WGA co-localization. Representative images of mTOR (red), VPS34 (green) and WGA (blue) stains at each time point are provided (A). Quantification of VPS34-WGA (B) and mTOR-VPS34 (C) co-localization is presented as Pearson’s correlation coefficient. Data in B and C are presented as mean±SEM. Ψ denotes a significant effect of time (p<0.05). *denotes a significant differences at this time point compared to PRE (p<0.05). Data analyzed on SPSS using Repeated Measures ANOVA with Holm-Bonferroni *post hoc* comparisons conducted on Microsoft Excel. All analyses – n=8.

### In vitro experiments

In C2C12 myotubes, a significant effect of treatment was found for S6K1^Thr389^ phosphorylation (p<0.001, Figure 5A). Here, nutrient/serum withdrawal significantly attenuated S6K1^Thr389^ phosphorylation compared to baseline levels (34% reduction, p<0.001), whereas phosphorylation was elevated by 65% and 50% in serum recovery (SR) and SR+SAR405 treatments respectively (both p<0.001, Figure 5A) with no difference between these two conditions (p=0.26). A treatment effect was also noted for 4EBP1^Thr37/46^ phosphorylation (p<0.001), however subsequent *post hoc* analysis revealed nutrient/serum withdrawal only significantly altered phosphorylation compared to baseline (∼28% reduction, p=0.015, Figure 5B). Again, no difference between SR and SR+SAR405 was observed (p=0.57).

**Figure 5.**
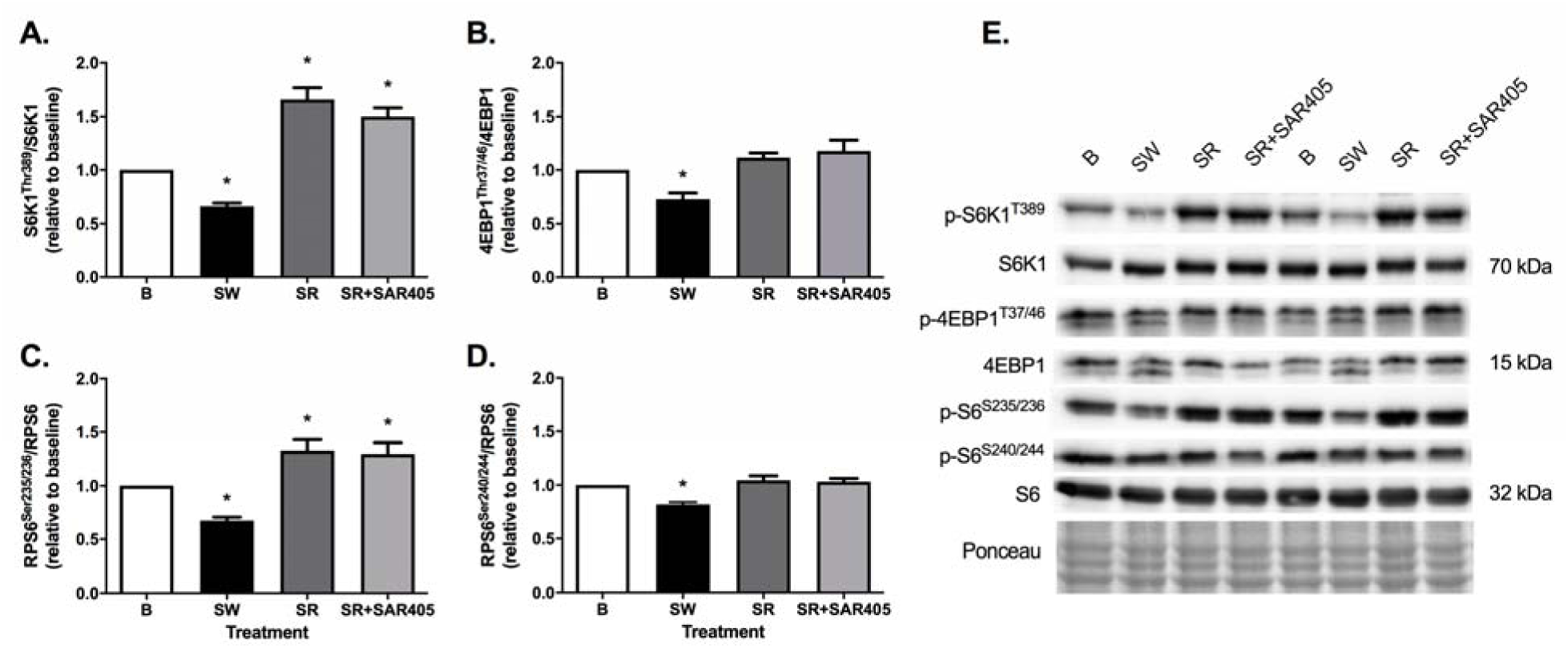
The effects of serum/nutrient withdrawal (∼14h) and subsequent serum recovery (30 min), +/- SAR405, on anabolic signalling in C2C12 myotubes (n=9/group). S6K1^Thr389^ (A), 4EBP1^Thr37/46^ (B), RPS6^Ser235/236^ (C) and RPS6^Ser240/244^ (D) phosphorylation were quantified in relation to their total proteins and ponceau staining was used as a loading control. Representative images are also provided (E). Data is presented in relation to baseline as Mean±SEM. *denotes a significant difference in this treatment compared to B (p<0.05). Data analyzed on SPSS using One-Way ANOVA with Holm-Bonferroni *post hoc* comparisons conducted on Microsoft Excel. All analyses – n=9. B = Baseline, SW = Serum Withdrawal & SR = Serum Recovery.

A significant treatment effect was also noted for both RPS6^Ser235/236^ and RPS6^Ser240/244^ phosphorylation (both p<0.001). RPS6^Ser235/236^ phosphorylation was significantly reduced by nutrient/serum withdrawal (33%, p<0.001, Figure 5C), whereas SR elicited a significant elevation in RPS6^ser235/236^ phosphorylation above baseline levels (32.7% increase, p=0.038, Figure 5C). A trend toward SR+SAR405 eliciting an elevation in RPS6^Ser235/236^ phosphorylation, compared to baseline, was also observed (29.5% increase, p=0.05) with no difference between the response of this treatment compared to SR (p=0.83). Following *post hoc* analysis of RPS6^Ser240/244^ phosphorylation, only serum/nutrient withdrawal altered phosphorylation status in relation to baseline (18% reduction, p<0.001, Figure 5D). No difference between SR and SR+SAR405 was observed (p=0.80). Representative immunoblots are displayed in Figure 5E.

In human primary myotubes, a significant treatment effect was noted for S6K1^Thr389^ phosphorylation (p<0.001). Here, serum/nutrient withdrawal reduced S6K1^Thr389^ phosphorylation by ∼70% compared to baseline (p=0.026, Figure 6A). SR and SR+SAR405 both elevated S6K1^Thr389^ phosphorylation above baseline levels (92% & 54%, p=0.026 & 0.035 respectively. Figure 6A). A trend for a greater response in SR, compared to SR+SAR405, was also observed (p=0.069). A treatment effect for 4EBP1^Thr37/46^ phosphorylation was also observed (p=0.004), however, following *post hoc* analysis no differences in 4EBP1^Thr37/46^ phosphorylation between individual treatment conditions was apparent (p>0.05, Figure 6B). Significant treatment effects were also observed for RPS6^Ser235/236^ and RPS6^Ser240/244^ phosphorylation, however *post hoc* analysis for both these variables did not reveal differences between individual treatments (p>0.05, Figures 6C & 6D respectively). Representative immunoblots are displayed in Figure 6E.

**Figure 6.**
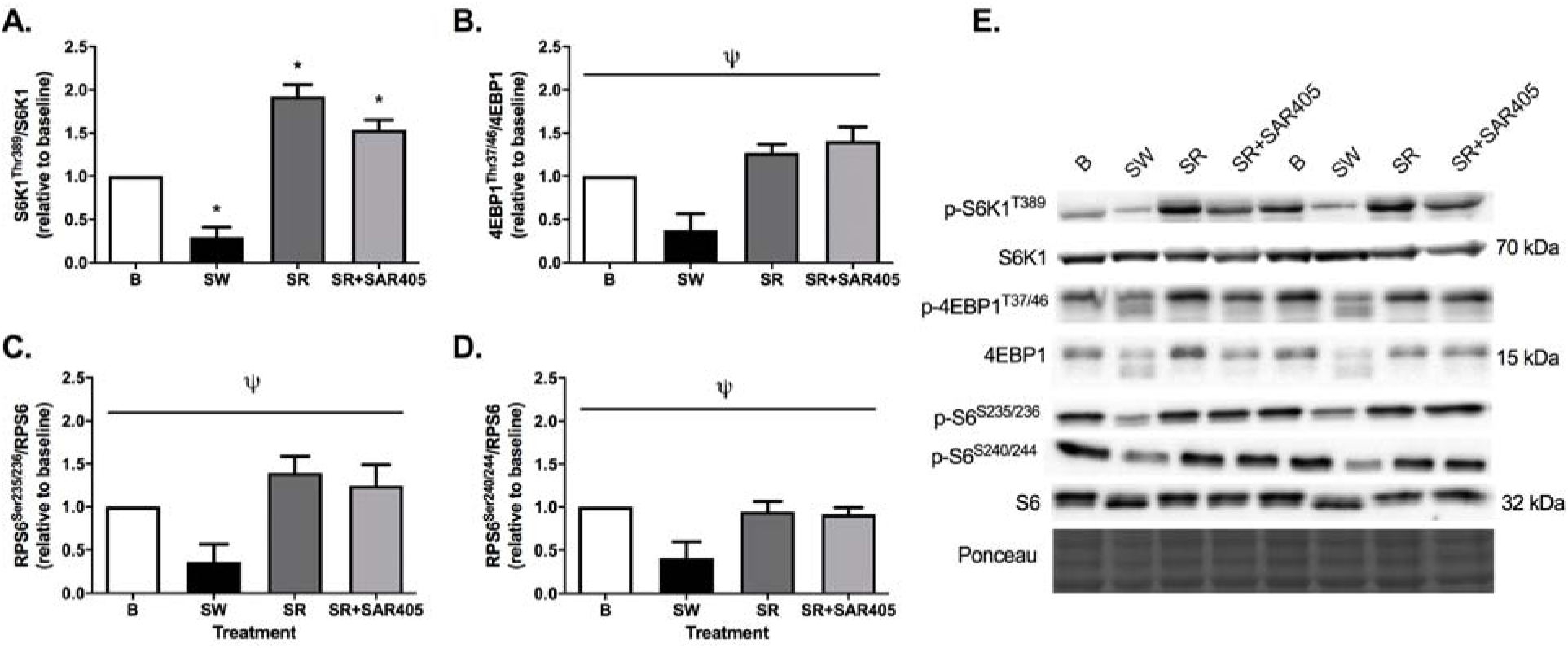
The effects of serum/nutrient withdrawal (∼14h) and subsequent serum recovery (30 min), +/- SAR405, on anabolic signalling in human primary myotubes (n=4). S6K1^Thr389^ (A), 4EBP1^Thr37/46^ (B), RPS6^Ser235/236^ (C) and RPS6^Ser240/244^ (D) phosphorylation were quantified in relation to their total proteins and ponceau staining was used as a loading control. Representative images are also provided (E). Data is presented in relation to baseline as Mean±SEM. *denotes a significant difference in this treatment compared to B (p<0.05). Ψ denotes a significant effect of treatment (p<0.05). Data analyzed on SPSS using Repeated Measures ANOVA with Holm-Bonferroni *post hoc* comparisons conducted on Microsoft Excel. All analyses – n=4. B = Baseline, SW = Serum Withdrawal & SR = Serum Recovery.

## Discussion

The class III PI3Kinase, Vps34, has been proposed as a nutrient/amino acid sensitive regulator of mTORC1 activity (4, 10, 22). To examine Vps34 action in human skeletal muscle, we examined changes in Vps34 activity and cellular localization following PRO-CHO ingestion *in vivo* and assessed the effect of Vps34 inhibition on anabolic responses to nutrient availability *in vitro* in C2C12 and human primary myotubes. We observed that PRO-CHO ingestion altered Vps34 localization, promoting translocation to the cell periphery and co-localization with mTORC1. Of note, these changes occurred independent of alterations in Vps34 kinase activity. In parallel, our *in vitro* studies demonstrated that the Vps34 specific inhibitor SAR405 did not affect nutrient stimulated activation of mTORC1. Together, these observations suggest a change in Vps34 cellular location, rather than an increase in kinase activity, may contribute to mTORC1 nutrient sensing in human skeletal muscle, however, loss of Vps34 activity does not prevent nutrient stimulation of mTORC1 *in vitro*.

The finding that PRO-CHO ingestion did not increase Vps34 kinase activity was contrary to our hypothesis and contrasts previous studies (4, 22). Previously, it has been shown that high-frequency electrical stimulation, a potent stimulator of mTORC1 activity, elevated Vps34 kinase activity in rodent skeletal muscle, a response suggested by the authors to be mediated by contraction-induced elevations in intracellular leucine (17). Given the increase in plasma leucine reported in the current study, we would expect our feeding protocol to result in similar increases in intramuscular leucine (1). In human skeletal muscle, there is only one previous study to have assessed Vps34 kinase activity (26). Here, sprint exercise combined with PRO-CHO ingestion did not alter kinase activity, whereas exercise in the fasted state elicited a trend toward elevated activity ∼1.5h following the final exercise bout. Importantly, in combination with our findings, this suggests that Vps34 kinase is not solely activated by leucine in human skeletal muscle and may suggest that a contraction stimulus is needed to activate this kinase.

In an attempt to further clarify the role of Vps34 in mTORC1 activation in skeletal muscle, we completed *in vitro* experiments in both C2C12 and human primary myotubes, utilising the Vps34 specific inhibitor SAR405 (25). In support of our findings *in vivo*, we observed no effect of SAR405 administration on mTORC1 signaling responses to serum recovery in C2C12 or human primary myotubes, suggesting Vps34 kinase activity is not necessary for mTORC1 activation.

Recent work from our lab (11, 28), and others (15) suggests that mTORC1 activation in skeletal muscle involves the translocation of mTORC1-lysosome complexes to peripheral regions of the cell (12). Here, we report a similar process by which mTOR-LAMP2 co-localize in the fasted state, prior to mTOR-LAMP2 complex translocation post PRO-CHO ingestion. Vps34 has previously been implicated in mTOR translocation *in vitro*, where it is required for the recruitment of mTOR to lamellipodia (cellular projections of motile cells) in response to insulin stimulation, co-localizing with mTOR in these regions (10). In the current study, we also found Vps34 translocation toward the cell periphery following nutrient provision, with a trend toward a time effect noted for Vps34-WGA co-localization (p=0.053). In this context, Vps34-WGA co-localization increased significantly above basal fasted levels 1 hour post-feeding (p=0.043) before returning to basal fasted levels 3 hours post PRO-CHO ingestion. Therefore, our observation that Vps34 translocation, and localization with mTORC1, occurs in human skeletal muscle indicates that Vps34 may act as a scaffold for mTORC1 recruitment toward the cell periphery, with an increase in Vps34 kinase activity not required for this process.

From the current data it is not possible to conclude whether Vps34 and mTORC1 translocate in tandem or independently before co-localizing, or the physiological relevance of these events in human skeletal muscle. A potential mechanism as to how Vps34 may regulate mTORC1 translocation and activation has recently been proposed by Hong and colleagues (13) who suggested that the product of Vps34 kinase activity, PI(3)P, may regulate lysosomal positioning via its receptor, FYCO1 (13). In this model, AAs increase the association between FYCO1 and lysosomes, whereas the ablation of this protein caused the clustering of mTOR-positive lysosomes to perinuclear regions and attenuated mTORC1 activity irrespective of nutrient availability (13). Other potential mechanisms as to how Vps34 may regulate mTORC1 activity include via Tuberous Sclerosis Complex 2 (TSC2) ubiquitination (21) and leucyl t-RNA synthetase (LRS)-regulated mTORC1 activation (34), however each of these processes require further investigation to determine their relevance for mTORC1 activity in skeletal muscle.

In conclusion, we report that PRO-CHO ingestion does not increase Vps34 activity in human skeletal muscle, whilst pharmacological inhibition of Vps34 does not prevent nutrient stimulation of mTORC1. However, PRO-CHO ingestion did promote Vps34 translocation to the cell periphery, where Vps34/mTOR co-localize. Therefore, our data suggests that cellular trafficking of Vps34 may result from increased PRO-CHO availability and occur in order to increase Vps34 association with mTOR. Future research studying the effects of resistance exercise, independently or in combination with AA ingestion may be required to fully understand the role of Vps34 in nutrient sensing and skeletal muscle anabolism.

## Authors Contributions

N.H. & A.P. conceived the study. J.R.D. Z.S. L.B. & A.P. designed and conducted *in vivo* experiments. N.H. conducted and completed analysis for all *in vitro* experiments. N.H. J.R.D. Z.S. S.J. D.L.H. J.T.M. M.F.O. T.N. & S.W.J. performed analysis. N.H. completed data processing and statistical analysis. N.H. J.R.D. & A.P. drafted the manuscript. All authors approved the final version.

